# Age Grading *An. Gambiae* and *An. Arabiensis* Using Near Infrared Spectra and Artificial Neural Networks

**DOI:** 10.1101/490326

**Authors:** Masabho P. Milali, Maggy T. Sikulu-Lord, Samson S. Kiware, Floyd Dowell, George F. Corliss, Richard J. Povinelli

## Abstract

**Background:** Near infrared spectroscopy (NIRS) is currently complementing techniques to age-grade mosquitoes. NIRS classifies lab-reared and semi-field raised mosquitoes into < or ≥ 7 days old with an average accuracy of 80%, achieved by training a regression model using partial least squares (PLS) and interpreted as a binary classifier.

**Methods and findings:** We explore whether using an artificial neural network (ANN) analysis instead of PLS regression improves the current accuracy of NIRS models for age-grading malaria transmitting mosquitoes. We also explore if directly training a binary classifier instead of training a regression model and interpreting it as a binary classifier improves the accuracy.

A total of 786 and 870 NIR spectra collected from laboratory reared *An. gambiae* and *An. arabiensis*, respectively, were used and pre-processed according to previously published protocols. Based on ten-fold Monte Carlo cross-validation, an ANN regression model scored root mean squared error (RMSE) of 1.6 ± 0.2 for *An. gambiae* and 2.8 ± 0.2 for *An. arabiensis*; whereas the PLS regression model scored RMSE of 3.7 ± 0.2 for *An. gambiae*, and 4.5 ± 0.1 for *An. arabiensis*. When we interpreted regression models as binary classifiers, the accuracy of the ANN regression model was 93.7 ± 1.0 % for *An. gambiae*, and 90.2 ± 1.7 % for *An. arabiensis*; while PLS regression model scored the accuracy of 83.9 ± 2.3% for *An. gambiae*, and 80.3 ± 2.1% for *An. arabiensis*. We also find that a directly trained binary classifier yields higher age estimation accuracy than a regression model interpreted as a binary classifier. A directly trained ANN binary classifier scored an accuracy of 99.4 ± 1.0 for *An. gambiae*, and 99.0 ± 0.6% for *An. arabiensis*; while a directly trained PLS binary classifier scored 93.6 ± 1.2% for *An. gambiae*, and 88.7 ± 1.1% for *An. arabiensis*.

**Conclusion:** Training both regression and binary classification age models using ANNs yields models with higher estimation accuracies than when the same age models are trained using PLS. Regardless of the model architecture, directly trained binary classifiers score higher accuracy on classifying age of mosquitoes than a regression model translated as binary classifier. Therefore, we recommend training models to estimate age of *An. gambiae* and *An. arabiensis* using ANN model architectures and direct training of binary classifier instead of training a regression model and interpret it as a binary classifier.

## Introduction

Estimating the age of mosquitoes is one of the indicators used by entomologists for estimating vectorial capacity [1] and the effectiveness of an existing mosquito control intervention. Malaria is a vector-borne parasitic disease transmitted to people by mosquitoes of the genus *Anopheles*. The disease killed approximately 445,000 people in 2016 [2]. Mosquitoes contribute to malaria transmission by hosting and allowing the development to maturity of the malaria-causing *Plasmodium* parasite [3]. Depending on environmental temperature, *Plasmodium* takes 10-14 days in an *Anopheles* mosquito to develop fully enough to be transmitted to humans [3]. Therefore, knowing the age of a mosquito provides an indication of whether a mosquito is capable of transmitting malaria.

Knowing the age of a mosquito population is also important when evaluating the effectiveness of a mosquito control intervention. Commonly used vector control interventions such as insecticide treated nets (ITNs) and indoor residual spraying (IRS) reduce the abundance and the lifespan of a mosquito population to a level that does not support *Plasmodium* parasite development to maturity [4, 5]. Monitoring and evaluation of ITNs and IRS involves determining the age and species composition of the mosquito population before and after intervention. The presence of a small number of old mosquitoes in an area with an (ITNs or IRS) intervention indicates that the intervention is working. On the other hand, if there are more old mosquitoes, the intervention is not working effectively.

The current techniques used to estimate mosquito age are based on a combination of ovary dissecting and conventional microscopy to determine their egg laying history. Those found to have laid eggs are assumed to be older than those found to not have laid eggs [6]. This assumption can be misleading, as mosquitoes can be old but have not laid eggs and can be young (at least three days old), and have laid eggs. Dissection is laborious, difficult, and limited to only few experts. As a result, we need a new approach to address these limitations.

Different techniques such as a change in abundance of cuticular hydrocarbons [7, 8], transcriptional profiles [9, 10], and proteomics [11, 12] have been developed to age grade *Anopheles* mosquitoes. However, these techniques are still in early development stages and are limited to analyzing a small number of samples due to high analysis costs involved.

Near infrared spectroscopy (NIRS) is a complementary method to the current mosquito age grading techniques [13, 14]. NIRS is a high throughput technique, which measures the amount of the near infrared energy absorbed by samples. NIRS has been applied to identify species of insects infecting stored grains [15]; to age grade houseflies [16], stored-grain pests [17], and biting midges [18]; to differentiate between species and subspecies of termites [19]; to estimate the age and to identify species of morphologically indistinguishable laboratory reared and semi-field raised *Anopheles gambiae* and *Anopheles arabiensis* mosquitoes [13, 14, 20–23]; to estimate the age of *Aedes aegypti* mosquitoes [24]; and to detect and identify two strains of *Wolbachia pipientis* (wMelPop and wMel) in male and female laboratory-reared *Aedes aegypti* mosquitoes [25].

The current state-of-the-art of the accuracy of NIRS to classify the age of lab-reared *An. gambiae* and *An. arabiensis* is an average of 80% [13, 14, 20–23]. This accuracy is based on a trained regression model using partial least squares (PLS) and interpreted as a binary classifier to classify mosquitoes into two age groups (< 7 days and ≥ 7 days).

In this paper, using a set of spectra collected from lab-reared *An. gambiae* and *An. arabiensis*, we explored ways to improve the reported accuracy of a PLS model for estimating age of malaria-transmitting mosquitoes. Selection of a method to train a model is one of the important factors influencing the accuracy of the model [26]. Studies [27–30] compared the accuracies of artificial neural network (ANN) and PLS regression models for predicting respiratory ventilation; explored the application of ANN and PLS to predict the changes of anthocyanins, ascorbic acid, total phenols, flavonoids, and antioxidant activity during storage of red bayberry juice; determined glucose multivariation in whole blood using partial least-squares and artificial neural networks based on mid-infrared spectroscopy; and compared modeling of nonlinear systems with artificial neural networks and partial least squares, concluding that ANN models generally perform better than PLS models. Therefore, using ANN [29–31] and PLS, we trained regression age models and compared results.

Since previous studies [13, 14, 20–23] trained a regression model and interpreted it as a binary classifier (< 7 d and ≥ 7 d), the interpretation process may introduce errors and compromise the accuracy of the model. We further trained ANN and PLS binary classifiers and compared their accuracies with the ANN and PLS regression models translated as binary classifiers.

We find that training of both regression and binary classification models using an artificial neural network architectures yields higher accuracies than when the corresponding models are trained using partial least squares model architectures. Also, regardless of the architecture of the model, training a binary classifier yields higher age class estimation accuracy than a regression model interpreted as a binary classifier.

## Materials and methods

### Ethics approval

Permission for blood feeding laboratory-reared mosquitoes was obtained from the Ifakara Health Institute (IHI) Review Board, under Ethical clearance No. IHRDC/EC4/CL.N96/2004. Oral consent was obtained from each adult volunteer involved in the study. The volunteers were given the right to refuse to participate or to withdraw from the experiment at any time.

### Mosquito and spectra collection

We used spectra of *Anopheles gambiae* mosquito collected at 1, 3, 5, 7, 9, 11, 15, and 20 days and *An. arabiensis* collected at 1, 3, 5, 7, 9, 11, 15, 20 and 25 days post emergence from the Ifakara Health Institute insectary. While *An. arabiensis* were reared in a semi-field system (SFS) at ambient conditions, *An. gambiae* were reared in a room made of bricks at controlled conditions. Adult mosquitoes were often provided with a human blood meal in a week and 10% glucose solution daily. Using a LabSpec 5000 NIR spectrometer with an integrated light source (ASD Inc., Longmont, CO), we followed the protocol supplied by Mayagaya and colleagues to collect spectra [13]. Prior to spectra collection, as opposed to killing by chloroform, mosquitoes were killed by freezing for 20 minutes. A total of 786 *An. gambiae* and 870 *An. arabiensis* were scanned with at least 70 mosquitoes from each age group.

### Model training

We first trained ANN and PLS regression models, scored and compared their accuracies as regressors and then as binary classifiers. We further trained binary classifiers and compared the accuracies with regressors interpreted as binary classifiers. We used a two-tail t-test to test the hypothesis that there is significant difference in accuracies between ANN and PLS trained model, a one-tail t-test to test the hypothesis that an ANN trained model scores higher accuracies than a PLS trained model.

In each species, we separately processed spectra according to Mayagaya et. al, randomized, and divided processed spectra into two groups. The first group contained 70% of the total spectra and was used for training models. The second group had 30% of the total spectra and was used for out-of-sample testing.

We trained a PLS ten-components model on using ten-fold cross validation [32]. Even though a range of six to ten PLS components were used in previous studies [13, 14, 20–22], we used ten PLS components after plotting the percentage of variance explained in the dependent variable against the number of PLS components (S1 Fig in the supporting information). For both species, there is not much change in the percentage variance explained in the dependent variables beyond ten components.

For the ANN model, we trained a feed-forward ANN with one hidden layer, ten neurons, and a linear transfer function (purelin) using Levenberg-Marquardt (damped least-squares) optimization [33]. We used actual mosquito ages as labels during training of both PLS and ANN regression models. We determined whether the trained models are over-fit by applying trained models (PLS and ANN) to estimate ages of mosquitoes on both training (in sample) and test (out-of-sample) data sets. Normally, if the model is not over-fit, the accuracy of the model is consistent between training and test sets [34].

The accuracies of the models were determined by computing their root mean squared error (RMSE) [35–37]. We evaluated the influence of the model architecture on the model accuracy by comparing their accuracies.

When interpreting the regression models as binary classifiers, mosquitoes with an estimated age < 7 days were considered as less than seven days old, and those ≥ 7 were considered older than or equal to seven days old. Using Equations 1, 2, and 3, we computed and compared sensitivity, specificity, and accuracy between the PLS and ANN regression models interpreted as binary classifiers. Sensitivity of the model is the ability to classify mosquitoes correctly, which are older than or equal to seven days old (assumed to be positively related to malaria transmission), and specificity is the ability of the model to classify mosquitoes correctly which are less than seven days old (assumed to be negatively related to malaria transmission) [38–40].

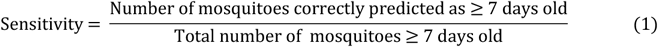

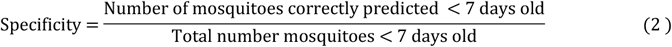

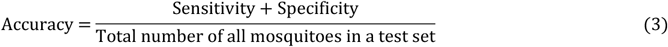

Training a regression model and interpreting it as a binary classifier can compromise the accuracy of the model as a classifier. This is because, while training a regression model forces the model to learn differences between actual ages of mosquitoes, direct training of a binary classifier forces the model to learn similarities between mosquitoes of the same class and only differences between two classes. Therefore, we directly trained binary classification models using ANN and PLS architectures and compare the accuracies with the ANN and PLS regression models interpreted as binary classifiers. In both species, we divided processed spectra (786 spectra for *An. gambiae* and 870 spectra for *An. arabiensis*) into two groups; < 7 days old and ≥ 7 days old. The spectra in a group with mosquitoes < 7 days old were labeled 0, 1 for those in a group with mosquitoes ≥ 7 days old, and the two groups were merged. The spectra were randomized and divided into training (N = 508 for both species) and test (N = 278 for *An. gambiae* and N = 362 for *An. arabiensis*) sets. We trained a PLS ten-component model using ten-fold cross-validation [32] and a one hidden layer, ten neuron feed-forward ANN using logistic regression as a transfer function and Levenberg-Marquardt (damped least-squares) optimization for training [33, 41]. During interpretation of these models, mosquitoes < 0.5 were considered as < 7 days old and ≥ 0.5 as ≥ 7 days old. Using Equations 1, 2, and 3, for each species, we computed specificity, sensitivity, and accuracy of the trained PLS and ANN binary classifiers and compared to the PLS and ANN regressors interpreted as the binary classifiers.

We repeated the process of random splitting the dataset into training and test sets; training, testing and scoring the accuracies of trained models ten times and compare the average results, a process known as Monte Carlo cross-validation [42–44].

## Results

Both PLS and ANN regression models consistently estimated the age of *An. gambiae* and *An. arabiensis* in the training and test data sets, showing that the models were not over-fit during training (Fig 1). Table 1, S2 Fig in the supporting information and Fig 2 present the performances of PLS and ANN regression models when estimating actual age of *An. gambiae* and *An. arabiensis* in the test data set and when their outputs are interpreted into two age classes, showing significant differences in accuracies of the two models (PLS vs ANN models). Table 1 further shows that the ANN regression model scores significantly higher accuracy than the PLS regression model.

**Fig 1:**
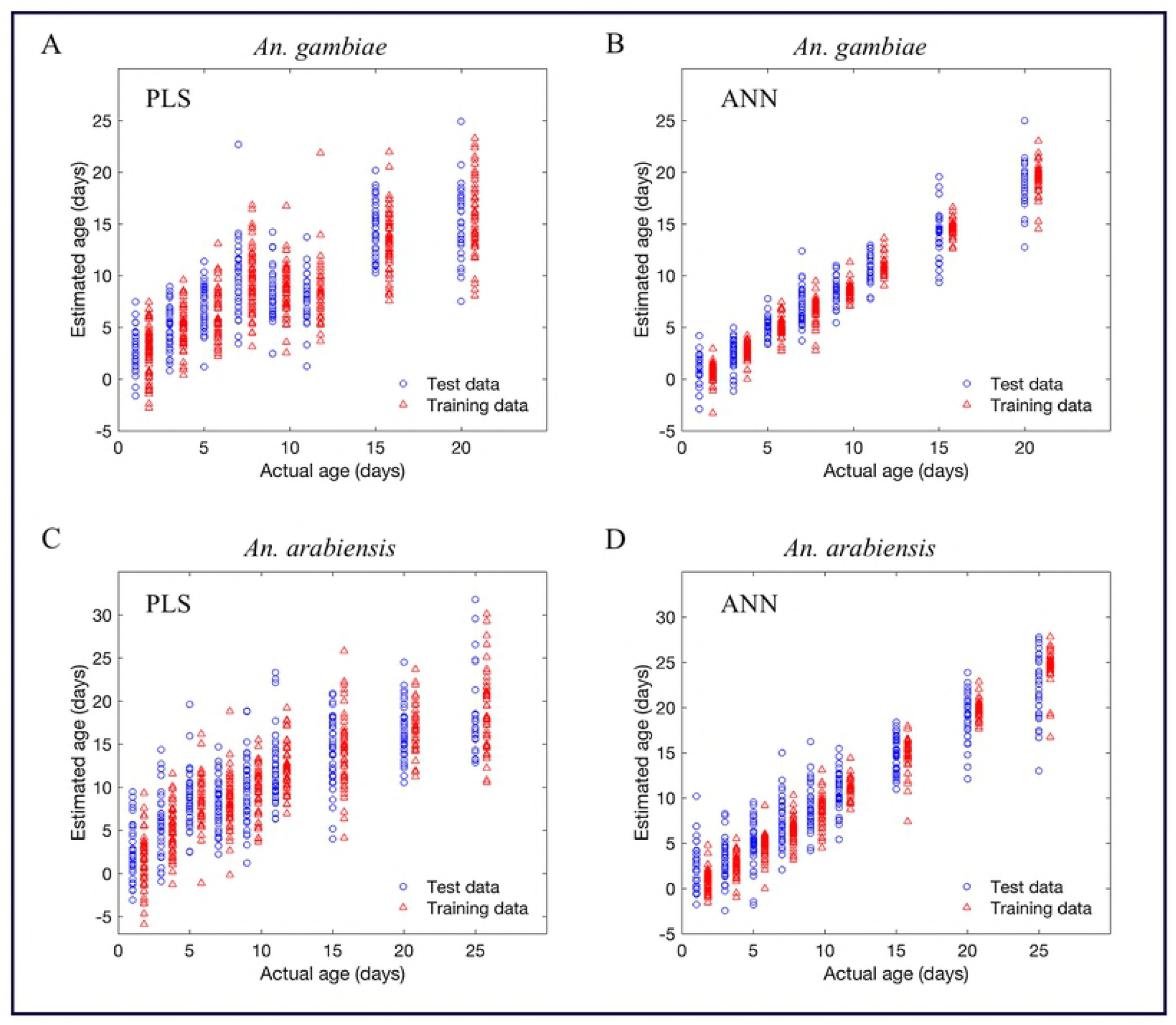
PLS (A and C) and ANN (B and D) regression models, estimating actual age of training and testing samples of *An. gambiae* (A and B) and *An. arabiensis* (C and D), respectively.

**Table 1:** Performance analysis of PLS and ANN regression models on estimating age of *An. gambiae* and *An. arabiensis*. Results from ten-fold Monte Carlo cross-validation.

| Species | Model estimation | Metric | Model architecture |  | P-value<br>(two tail) | P-value<br>(one tail) |
| --- | --- | --- | --- | --- | --- | --- |
|  |  |  | PLS | ANN |  |  |
| *An. gambiae* | Actual age | RMSE | 3.7 ± 0.2 | 1.6 ± 0.2 | 3.9 x 10<sup>-09</sup> | 1.6 x 10<sup>-11</sup> |
|  | Age class | Accuracy (%) | 83.9 ± 2.3 | 93.7 ± 1.0 | 3.6 x 10<sup>-07</sup> | 2.3 x 10<sup>-07</sup> |
|  |  | Sensitivity (%) | 89.0 ± 2.1 | 92.5 ± 1.6 | 4.7 x 10<sup>-02</sup> | 4.7 x 10<sup>-01</sup> |
|  |  | Specificity (%) | 75.8 ± 5.2 | 95.6 ± 1.8 | 3.7 x 10<sup>-11</sup> | 1.1 x 10<sup>-06</sup> |
| *An. arabiensis* | Actual age | RMSE | 4.5 ± 0.1 | 2.8 ± 0.2 | 1.7 x 10<sup>-09</sup> | 5.9 x 10<sup>-08</sup> |
|  | Age class | Accuracy (%) | 80.3 ± 2.1 | 90.2 ± 1.7 | 1.4 x 10<sup>-07</sup> | 2.4 x 10<sup>-08</sup> |
|  |  | Sensitivity (%) | 90.5 ± 1.9 | 91.7 ± 3.3 | 5.8 x 10<sup>-01</sup> | 6.0 x 10<sup>-01</sup> |
|  |  | Specificity (%) | 60.3 ± 4.2 | 88.4 ± 3.9 | 1.7 x 10<sup>-07</sup> | 1.2 x 10<sup>-06</sup> |

**Fig 2:**
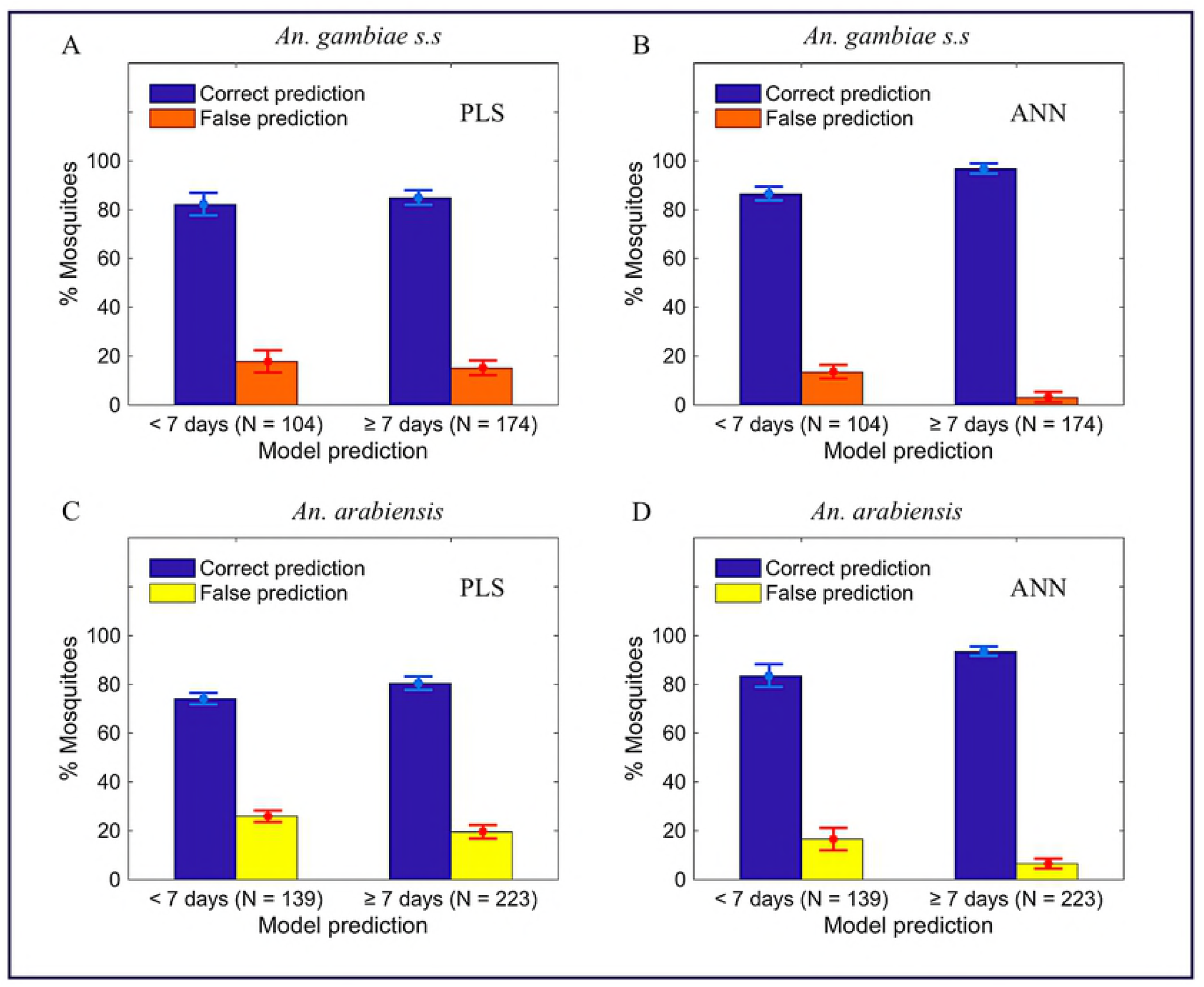
Number of *An. gambiae s.s* (A and B) and *An. arabiensis* (C and D) in two age classes (less than or greater/equal seven days) when PLS (A and C) and ANN (B and D) regression models, respectively, interpreted as binary classifiers.

Fig 3 represents consistency in accuracy of PLS (A and C) and ANN (B and D) directly trained binary classifiers on estimating both training and test data sets, showing that the models were not over-fitted during training. Fig 4, S3 Fig in the supporting information and Table 2 present the results when directly trained PLS (A and C) and ANN (B and D) binary classifiers were applied to classify ages of *An. gambiae* (A and B) and *An. arabiensis* (C and D) in test sets (out-of-the sample testing), showing ANN binary classifier scores higher accuracy than the PLS binary classifier. The results further show that in both species, irrespective of the architecture used to train the model, direct training of the binary classifier scores significantly higher accuracy, specificity, and sensitivity than the regression model translated as a binary classifier (S1 Table in the supporting information).

**Fig 3:**
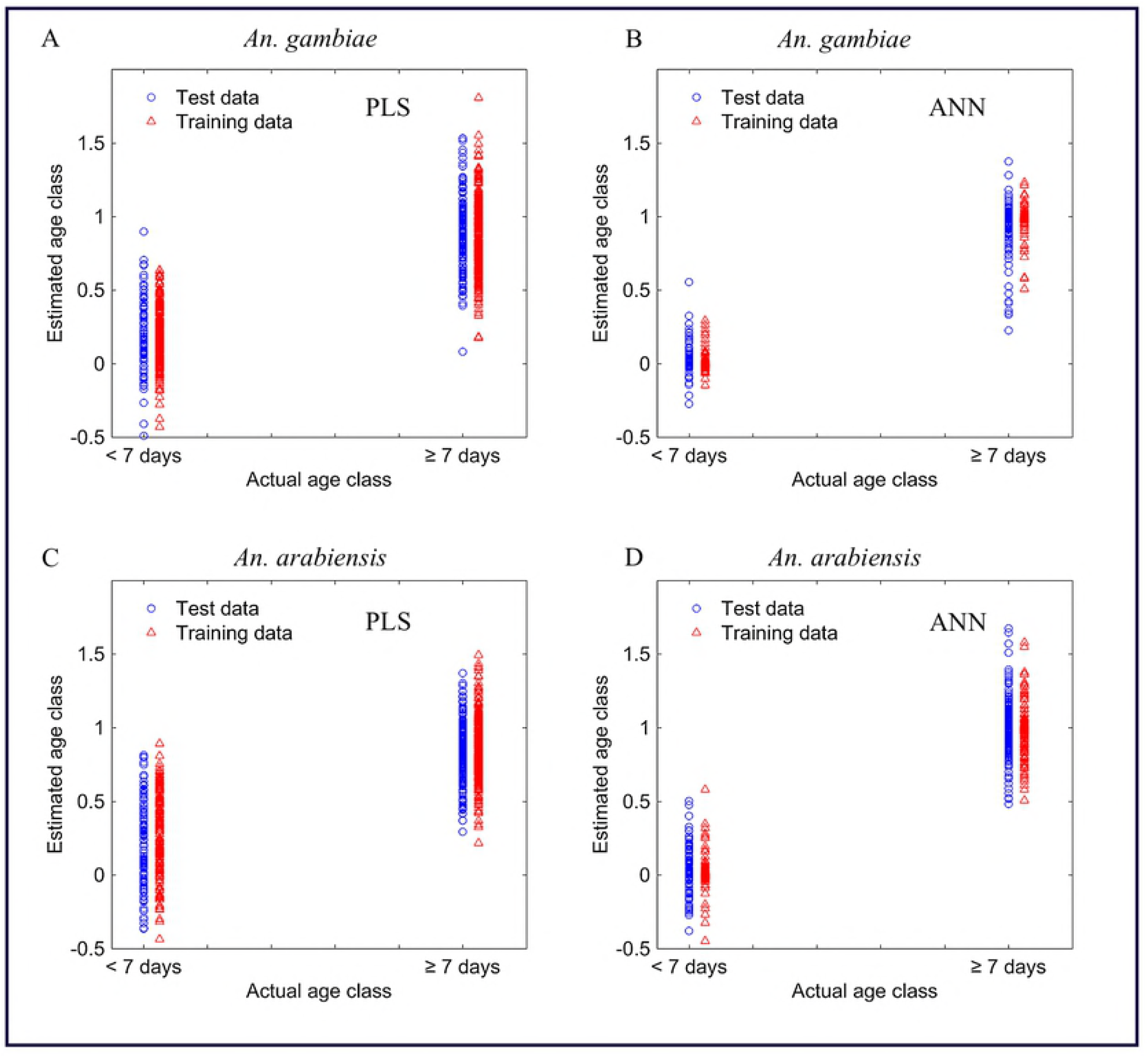
The consistency in accuracies of directly trained PLS (A and C) and ANN (B and D) binary classifiers for estimating age classes of *An.gambiae* (A and B) *and An. arabiensis* (C and D) in both training and testing sets.

**Fig 4:**
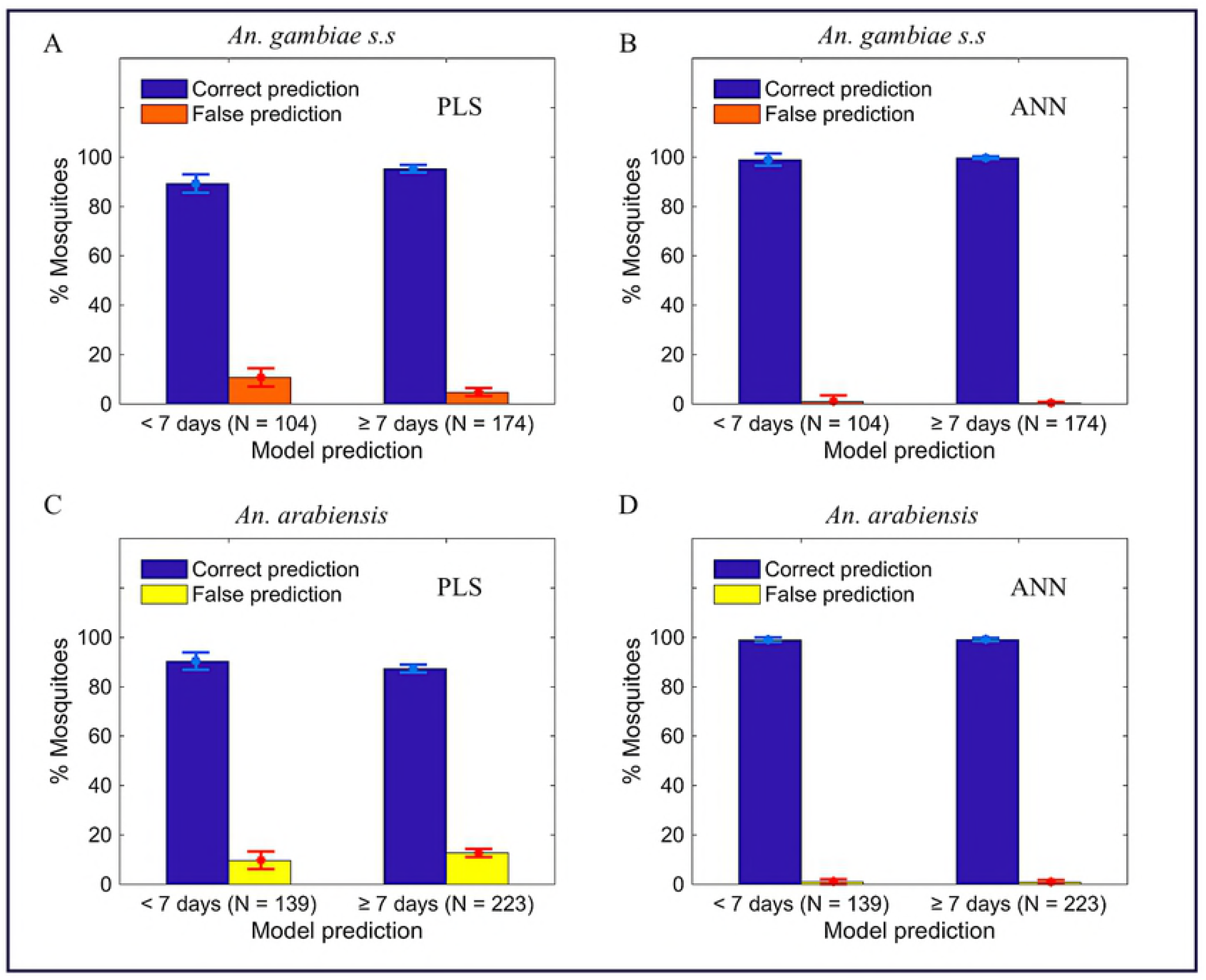
The number of correct and false predictions in each estimated age-class when directly trained PLS (A and C) and ANN (B and D) binary classifiers were applied to classify age of *An. gambiae* (A and B) and *An. arabiensis* (C and D) in testing sets. Results from ten replicates.

**Table 2:** Comparison of the accuracy of ANN and PLS classification models on ten replicates

| Species | Metric | Model architecture |  | P-value<br>(two-tail) | P-value<br>(one-tail) |
| --- | --- | --- | --- | --- | --- |
|  |  | PLS | ANN |  |  |
| *An. gambiae* | Accuracy (%) | 93.6 ± 1.2 | 99.4 ± 1.0 | 2.4 x 10<sup>-19</sup> | 1.2 x 10<sup>-19</sup> |
|  | Sensitivity (%) | 94.4 ± 1.6 | 99.3 ± 1.4 | 1.6 x 10<sup>-04</sup> | 2.0 x 10<sup>-05</sup> |
|  | Specificity (%) | 92.4 ± 1.9 | 99.5 ± 0.7 | 2.2 x 10<sup>-06</sup> | 6.0 x 10<sup>-05</sup> |
| *An. arabiensis* | Accuracy (%) | 88.7 ± 1.1 | 99.0 ± 0.6 | 1.5 x 10<sup>-21</sup> | 7.6 x 10<sup>-22</sup> |
|  | Sensitivity (%) | 95.4 ± 1.4 | 99.5 ± 0.5 | 4.5 x 10<sup>-05</sup> | 2.3 x 10<sup>-05</sup> |
|  | Specificity (%) | 75.2 ± 3.4 | 98.3 ± 1.3 | 4.0 x 10<sup>-09</sup> | 2.0 x 10<sup>-09</sup> |

## Discussion

This study aimed at improving the current state of the art accuracies of the models trained using near infrared spectra to estimate the age of *An. gambiae* and *An. arabiensis*. Previous studies [13, 14, 20–23] trained a regression model using partial least squares (PLS) and interpreted it as a binary classifier (< 7 d and ≥ 7 d) with an accuracy around 80%.

Knowing that the selection of a model architecture often influences the model accuracy [26], we trained age regression models using an artificial neural network [29–31, 45, 46] and partial least squares as model architectures and compared the accuracies. ANN models achieved significantly higher accuracies than corresponding PLS regression models. As summarized in Table 1, ANN regression models scored an average RMSE of 1.60 ± 0.18 for *An. gambiae* and 2.81 ± 0.22 for *An. arabiensis*. The PLS regression models scored RMSE of 3.66 ± 0.23 for *An. gambiae* and 4.49 ± 0.09 for *An. arabiensis*. When both ANN and PLS regression models were interpreted as binary classifiers, ANN regression model scored accuracy, sensitivity, and specificity of 93.71 ± 1.03%, 92.54 ± 1.60%, and 95.64 ± 1.82%, respectively, for *An. gambiae;* 90.16 ± 1.70%, 91.68 ± 3.27% and 88.44 ± 3.86%, respectively, for *An. arabiensis*. The PLS regression model scored accuracy, sensitivity, and specificity of 83.85 ± 2.32%, 89.00 ± 2.10%, and 75.82 ± 5.22%, respectively, for *An. gambiae;* 80.30 ± 2.06%, 90.48 ± 1.88%, and 60.25 ± 4.20%, respectively, for *An. arabiensis*.

The interpretation of a regression model into a binary classifier can introduce errors that compromise the accuracy of the model. We directly trained PLS and ANN binary classifiers and compared the accuracies with ANN and PLS regression models interpreted as binary classifiers. Irrespective of the model architecture, directly trained binary classifiers scored significantly higher accuracies than corresponding regression models interpreted as binary classifiers (S1 Table in the supporting information). The explanation of these results could be that, training a regression model and interpreting it as a binary classifier involved learning differences between multiple age groups (1, 3, 5, 7, 9, 11, 13, 15, and 20 days old for *An. gambiae* and 1, 3, 5, 7, 9, 11, 13, 15, 20 and 25 days for *An. arabiensis)* of mosquitoes, which can be challenging for two consecutive age groups. In contrast, direct training of the binary classifier involved learning differences existing between only two age groups. During direct training of the binary classifier, the process of dividing spectra into two groups (< 7 or ≥ 7 days) forced a model to learn similarities instead of differences between mosquitoes of the same age class. We also observed that directly trained ANN binary classifier scored higher accuracy than directly trained PLS binary classifier. ANN binary classifier scored an accuracy, sensitivity, and specificity of 99.4 ± 1.0%, 99.3 ± 1.4%, and 99.5 ± 0.7%, respectively, for *An. gambiae;* 99.0 ± 0.6%, 99.5 ± 0.5%, and 98.3 ± 1.3%, respectively, for *An. arabiensis*. The PLS binary classifier scored 93.6 ± 1.2%, 94.4 ± 1.6%, and 92.5 ± 1.9% for *An. gambiae;* 88.7 ± 1.1%, 95.5 ± 1.4%, and 75.2 ± 3.5% for *An. arabiensis* (Table 2).

Our study is not the first to observe ANN model outperforming PLS model. These findings are supported with other previous studies [27–29, 31] compared the accuracies of ANN and PLS models, where they report ANN perform better than PLS. The explanation on these results could be that ANN, unlike PLS, considers both linear and unknown non-linear relationships between dependent and independent variables [29–31]; builds independent-dependent relationships that interpolates well even to cases that were not exactly presented by training data; and has a self mechanism of filtering and handling noisy data during training [45, 46]. Hence, ANN models are unbiased estimators in contrast to PLS models (Fig 5 and S4 Fig in supporting information).

**Fig 5:**
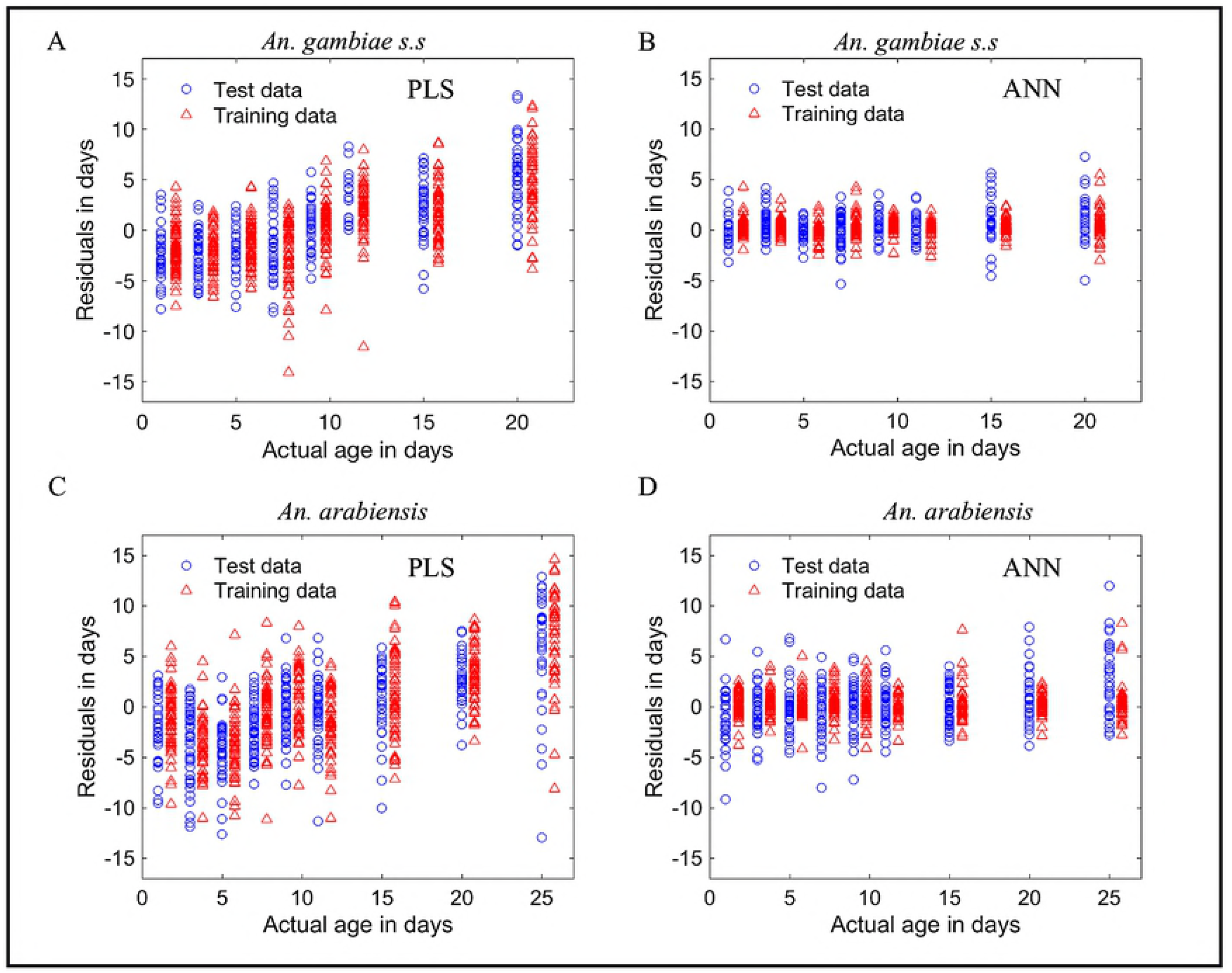
Error distribution per actual age of *An. gambiae* and *An. arabiensis* when ANN and PLS regressors applied to estimate the actual ages of mosquitoes in training and test data sets, showing uniform distribution of errors (un-biased estimating) across actual ages of mosquitoes for ANN regressor and un-uniform distribution of errors (biased estimating) for PLS regressor.

## Conclusion

We conclude that training both regression and binary classification age artificial neural network models yield higher accuracies than partial least squares models. Also, training a binary classifier scores higher accuracy than training a regression model and interpreting it as a binary classifier. Hence, we recommend training of age models using artificial neural network and training of binary classifier instead of training regression model and interpret it as binary classifier.

## Acknowledgements

We thank Andrew Kafwenji and Paulina Kasanga for help maintaining the mosquito colony, Marta F. Maia, Fredros O. Okumu, and Sheila Ogoma for participating in grant writing and managing of the project produced the data used in this manuscript, and the USDA, Agricultural Research Service, Center for Grain and Animal Health Research, USA for loaning us the near-infrared spectrometer used to scan the mosquitoes. We also thank Michael Henry and Nikita Lysenko for helping with mosquito scanning to collect spectra in Tanzania. Finally, Gustav Mkandawile who worked tirelessly to make sure we obtained mosquitoes and Ben Durette who participated in the initial stages of data analysis.

Mention of trade names or commercial products in this publication is solely for the purpose of providing specific information and does not imply recommendation or endorsement by the U.S. Department of Agriculture. USDA is an equal opportunity provider and employer.

## Supporting information

**S1 Fig. The percentage of variance explained in the dependent variable against the number of PLS components:** A) *An. gambiae* B) *An. arabiensis* (TIFF)

**S2 Fig. Box plot showing the performance of ANN and PLS regression models when applied to estimate the actual ages of *An. gambiae* and *An. arabiensis* in the test data set. (TIFF)**

**S3 Fig. Box plot showing the performance of a directly trained ANN and PLS binary classifiers when applied to estimate the age classes of *An. gambiae* and *An. arabiensis* in the test data set. (TIFF)**

**S4 Fig. Error distribution per actual age class of *An. gambiae* and *An. arabiensis* when directly trained ANN and PLS binary classifiers applied to estimate age classes of mosquitoes in training and test data sets, showing uniform distribution of errors (un-biased estimating) across actual age classes of mosquitoes for ANN binary classifiers and un-uniform (biased estimating) distribution for PLS classifiers. (TIFF)**

**S1 Table: Comparison of accuracies between directly trained binary classifiers and regressers interpreted as binary classifiers. Results from ten-fold Monte Carlo cross-validation (TIFF and DOCX)**

**S1 Appendix. Excel file with the *Anopheles gambiae* spectra used in the analysis.** Column header, wavelengths in ‘nm’. (XLSX)

**S2 Appendix. Excel file with the *Anopheles arabiensis* spectra used in the analysis.** Column header, wavelengths in ‘nm’. (XLSX)

**S3 Appendix. Matlab code used to run the analysis for *Anopheles gambiae*. (M)**

**S4 Appendix. Matlab code used to run the analysis for *Anopheles arabiensis*. (M)**

